# Caveolae coupling of melanocytes signaling and mechanics is required for human skin pigmentation

**DOI:** 10.1101/666388

**Authors:** Lia Domingues, Ilse Hurbain, Floriane Gilles-Marsens, Nathalie André, Melissa Dewulf, Maryse Romao, Christine Viaris de Lesegno, Cédric Blouin, Christelle Guéré, Katell Vié, Graça Raposo, Christophe Lamaze, Cédric Delevoye

**Author notes:** Institut NeuroMyoGene, UCBL1, UMR 5310, INSERM U1217, Génétique et neurobiologie de C. Elegans, Faculté de Médecine et de Pharmacie, 8 Avenue Rockefeller, 69008 Lyon. these authors contributed equally (Graça Raposo, Christophe Lamaze).

## Abstract

Tissue homeostasis requires regulation of cell-cell communication, which relies on signaling molecules and cell contacts. In skin epidermis, keratinocytes secrete specific factors transduced by melanocytes into signaling cues to promote their pigmentation and dendrite outgrowth, while melanocytes transfer melanin pigments to keratinocytes to convey skin photoprotection. How epidermal cells integrate these functions remains poorly characterized. Here, we found that caveolae polarize in melanocytes and are particularly abundant at melanocyte-keratinocyte interface. Caveolae in melanocytes are sensitive to ultra-violet radiations and miRNAs released by keratinocytes. Preventing caveolae formation in melanocytes results in increased production of intracellular cAMP and melanin pigments, but decreases cell protrusions, cell-cell contacts, pigment transfer and epidermis pigmentation. Altogether, our data establish that, in melanocytes, caveolae serve as key molecular hubs that couple signaling outputs from keratinocytes to mechanical plasticity. This process is crucial to maintain cell-cell contacts and intercellular communication, skin pigmentation and tissue homeostasis.

## Introduction

Human skin comprises a highly stratified epidermis and a bottom dermis. The epidermis, the outermost and photo-protective layer of the skin, is mainly composed of melanocytes and keratinocytes that together create a structural and functional epidermal unit (Fitzpatrick and Breathnach, 1963). Melanocytes are neural crest-derived cells (Christiansen et al., 2000) that extend dendrites to contact up to 40 epidermal keratinocytes (Quevedo, 1972). The main role of melanocytes is to produce the melanin pigments in a specialized organelle, called melanosome, that undergoes maturation from early non-pigmented to late pigmented stages (Raposo and Marks, 2007). The maturing and pigmented melanosome moves towards the tip of the dendrites (Hume et al., 2001, 2007; Wu et al., 1998) to be transferred to keratinocytes where it protects the nuclei against ultra-violet (UV) radiations. In melanocytes, the formation of dendrites, melanosome biogenesis, and synthesis and transfer of melanin to keratinocytes is a tightly coordinated process under the control of UV radiations, keratinocytes-secreted factors and secreted endosomal-derived vesicles called exosomes (Abdel-Malek et al., 1994; Lo Cicero et al., 2015; Hirobe, 2005, 2014). From those, secreted hormones trigger different transduction pathways in melanocytes, including the cyclic adenosine monophosphate (cAMP) signaling pathway through binding to various G-protein coupled receptors (GPCRs) at the cell surface (D’Mello et al., 2016; Saldana-Caboverde and Kos, 2011). As a consequence, melanocytes increase pigment synthesis and dendrite outgrowth through regulation of Rho GTPases activity and remodeling of the actin cytoskeleton (Buscà and Ballotti, 2000; Buscà et al., 1998; Scott, 2002; Scott and Leopardi, 2003). We have recently shown that specific miRNAs associated with keratinocyte exosomes modulate human melanocyte pigmentation by enhancing the expression of proteins associated with melanosome maturation and trafficking (Lo Cicero et al., 2015). However, how environmental cues are spatially and temporally controlled in melanocytes to be efficiently translated into biochemical and physical cellular responses remains mostly uncharacterized.

Caveolae are cup-shaped plasma membrane invaginations firstly described in endothelial and epithelial cells (Palade, 1953; Yamada, 1955). Their size (50-100 nm) and the absence of an electron-dense coat morphologically distinguish caveolae from other invaginated structures at the plasma membrane (Stan, 2005). Caveolae are mainly composed of two groups of proteins, the caveolins (Cav1, 2 and 3) and the more recently identified cavins (Cavin1, 2, 3 and 4) (Bastiani et al., 2009; Hill et al., 2008; Kurzchalia et al., 1992; Liu et al., 2008; Nishimoto et al., 2002; Rothberg et al., 1992; Way and Parton, 1995). Caveolae biogenesis and functions are dependent on Cav1 and Cavin1 in non-muscle cells, and on Cav3 in muscle cells (Hansen and Nichols, 2010). Caveolae play various crucial functions including endocytosis, lipid homeostasis, signal transduction and, the most recently identified, mechanoprotection (Cheng and Nichols, 2016; Lamaze et al., 2017). As a transduction platform, caveolae control the production of second messengers, such as cAMP, through local confinement of different elements of this signaling cascade (Harvey and Calaghan, 2012). Cav1 and −3 contain a scaffolding domain (CSD) located in the N-terminal region suggested to interact with transmembrane adenylate cyclases (tmACs), to inhibit their activities and thus control intracellular cAMP levels (Toya et al., 1998). In cardiomyocytes, caveolae participate in the compartmentalization of intracellular cAMP which can regulate cell contractility in distal regions of the heart and, therefore, its function (Wright et al., 2014, 2018). The mechanoprotective role of caveolae is associated with the maintenance of plasma membrane integrity when both, cells and tissues, experience chronical mechanical stress (Cheng et al., 2015; Lo et al., 2015; Parton et al., 2017; Sinha et al., 2011). Caveolae were recently shown to couple mechanosensing with mechanosignaling in human muscle cells, a process impaired in caveolae-associated muscle dystrophies (Dewulf et al., 2019).

Epidermal melanocytes and keratinocytes are in constant communication, not only via secreted factors and exosomes that modulate cellular responses, but also by the physical contacts they establish to maintain the tissue homeostasis and pigmentation. Here, we report a new function for caveolae, which, by integrating the biochemical and mechanical behavior of melanocytes, control melanin transfer to keratinocytes and epidermis pigmentation. Altogether, this study provides the first evidence for a physiologic role of caveolae as a molecular sensing platform required for the homeostasis of the largest human tissue, the skin epidermis.

## Results

### Caveolae polarize in melanocytes and are positively-regulated by keratinocytes-secreted factors

Melanocytes and keratinocytes establish a complex intercellular dialogue required for skin photoprotection. 2D-co-culture systems, where these two cell types share the same medium, have been widely used to study intercellular communication and pigment transfer between epidermal cells (Hirobe, 2005; Lei et al., 2002). To evaluate the distribution of caveolae within the epidermal unit in 2D, normal human melanocytes and keratinocytes were co-cultured and labelled for the two constituents of caveolae, Cav1 or Cavin1. Immunofluorescence microscopy revealed that both Cav1 and Cavin1, and therefore caveolae, were asymmetrically distributed in melanocytes (**Figures 1A and B**), which were identified by the abundant staining of the premelanosome protein PMEL [hereafter referred as melanin, see Experimental Procedures; (Raposo et al., 2001)]. This polarization was not observed in keratinocytes.

Cells can break their symmetry in response to local external chemical and/or mechanical cues such as signaling molecules and/or cell-cell contacts, respectively (Altschuler et al., 2008; Goehring and Grill, 2013; Ladoux et al., 2016; Rappel and Edelstein-Keshet, 2017; Verkhovsky et al., 1999). However, in the absence of any type of spatial signaling, cell polarization can occur randomly and spontaneously (Wedlich-Soldner and Li, 2003). When grown alone in the absence of any pre-existent signaling cues, one third of melanocytes presented polarized caveolae, as shown by the asymmetric distribution of endogenous Cav1 and Cavin1 (**Figures 1C and S1A**). This polarization was restricted to caveolae as the distribution of <u>c</u>lathrin-<u>c</u>oated <u>p</u>its (CCPs; **Figure S1B**, red), the canonical plasma membrane invaginated-structures mediating endocytosis (Mayor et al., 2014) was even. Interestingly, the number of melanocytes showing caveolae asymmetrically distributed doubled when co-cultured with keratinocytes (Figures 1A and C), while co-culture with HeLa cells had no effect (**Figures 1C** and **S1A**). This shows that the intrinsic polarization of caveolae in melanocytes is specifically enhanced by keratinocytes, either by cell-cell contacts and/or by keratinocytes secreted factors. To address the role of extracellular factors in caveolae polarization, melanocytes were incubated with the medium recovered from a confluent culture of keratinocytes (referred as <u>c</u>onditioned <u>m</u>edium, CM). Under this condition, we observed a two-fold increase of the number of melanocytes with polarized caveolae as compared to cells grown in their own medium (**Figures 1D** and **S1C**). The proportion of melanocytes with polarized caveolae was similar between cells co-cultured with keratinocytes (**Figure 1C**) and cells incubated with conditioned medium (**Figure 1D**), which argues that factors secreted from keratinocytes are the main extracellular contributors to the increased polarization of caveolae in melanocytes.

**Figure 1.**
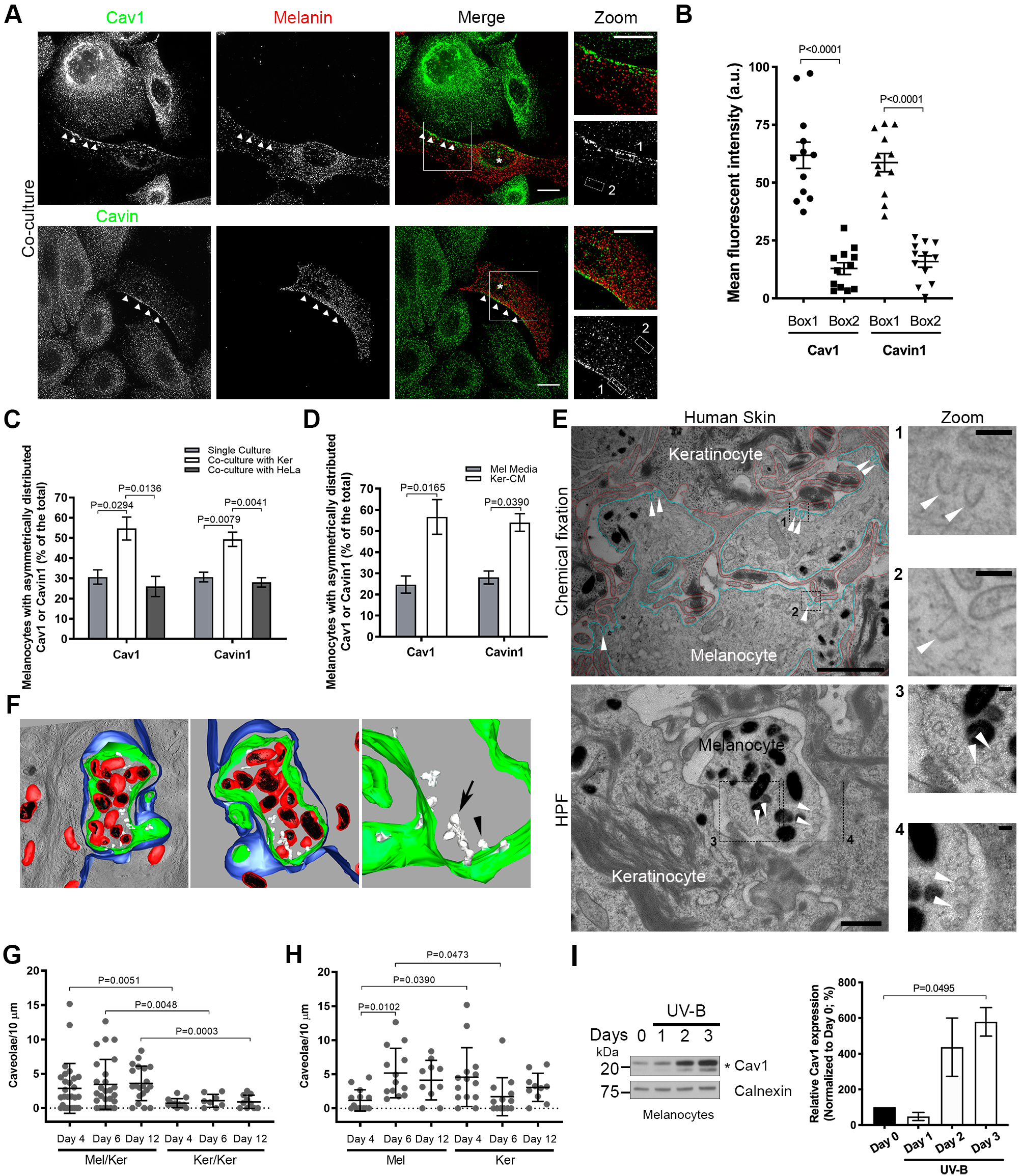
Caveolae localization and modulation in human epidermis and 2D co-culture. **A**. IFM images of melanocytes and keratinocytes co-cultured for 1 day, fixed, immunolabelled for Cav1 or Cavin1 (top or bottom, respectively; green) and melanin (HMB45, red). Arrowheads point Cav1 and Cavin1 polarization in melanocytes (white asterisks). The boxed regions mark the area zoomed in the insets. Bars, 10 µm. **B**. Quantification of Cav1 or Cavin1 mean fluorescent intensity in Boxes 1 and 2 depicted in the zoom panels A (n=12 cells). **C**. Quantification of the frequency of melanocytes displaying Cav1 or Cavin1 polarized (as in A, arrowheads) in mono- or co-culture with keratinocytes or HeLa (mono-culture, Cav1: 30.7 ± 3.5% and Cavin1: 30.7 ± 2.4%; co-culture with keratinocytes, Cav1: 54.7 ± 5.7% and Cavin1: 49.3 ± 3.5%; co-culture with HeLa cells, Cav1: 26.0 ± 5.0% and Cavin1: 28.0 ± 2.3%; n=150 cells, 3 independent experiments). **D**. Quantification of the frequency of melanocytes displaying Cav1 or Cavin1 polarized (as in A, arrowheads) after 14h incubation with supplemented medium or keratinocytes-conditional medium (Mel Medium, Cav1: 24.7 ± 4.1% and Cavin1: 28.0 ± 3.1%; Ker-CM, Cav1: 56.7 ± 8.2% and Cavin1: 54.0 ± 4.2%; n=150 cells, 3 independent experiments). **E**. Conventional 2D EM from human skin tissue fixed chemically (top) or immobilized by high pressure freezing (HPF, bottom). The plasma membranes of keratinocytes (red) and melanocytes (blue) were contoured manually (top). Arrowheads point plasma membrane invaginations with morphological features of caveolae. The boxed regions mark the area zoomed in the insets on the left. Bars: (main) 1 µm; (insets) 100 nm. **F**. 3D-model reconstruction by electron tomography of the melanocyte-keratinocyte interface at human skin epidermis; melanocytes plasma membrane (green), keratinocytes plasma membrane (blue), limiting membrane of pigmented melanosomes (red), melanin (black) and caveolae (white) in single (arrowhead) and clustered structures (arrow). See also **Video 1**.and **Figure S1F**. **G**. and **H**. Quantification during 3D-HRPE formation of the number of caveolae (as identified in E) per 10 µm of plasma membrane at the indicated interfaces (G) and of individual cell types at melanocyte-keratinocyte interface (H) (G, Mel-Ker: day 4, 2.9 ± 0.7, n=28; day 6, 3.4 ± 0.7, n=26; day 12, 3.6 ± 0.6, n=20; Ker-Ker: day 4, 0.7 ± 0.2, n=13; day 6, 1.1 ± 0.3, n=9; day 12, 0.9 ± 0.3, n=11; H, day 4, Mel: 1.2± 0.4, Ker: 4.5 ± 1.1; day 6, Mel: 5.0 ± 1.0, Ker: 1.7 ± 0.8; day 12, Mel: 4.1 ± 0.9; Ker: 3.1 ± 0.7; n= number of interfaces (G) or cells (H)). Note that 3D-HRPE stratifies at day 4, pigments at day 6 (normalized melanin content (a.u.) at day 4: 1, day 6: 2.46), and reaches completion at day 12. **I**. Immunoblot analysis and quantification of Cav1 protein levels in melanocytes exposed to daily radiations of U.V.-B (10 mJ/cm^2^) in three consecutive days (day 3; 573.3 ± 85.5%; n=2 independent experiments). Asterisk represents Cav1 full-length protein (upper band) and its truncated form (lower band). B-D and G-I, data are presented as mean ± s.e.m. B and D, paired t-test. G and H, comparison between interface/cells at the same time point: unpaired t-test with Welch’s correction; comparison between time points from the same cell type: one-way ANOVA with Tukey’s multiple-comparison test.

### Caveolae localize at the melanocyte-keratinocyte interface in human epidermis and accumulate in melanocytes during tissue pigmentation

We investigated the distribution of caveolae at the melanocyte-keratinocyte interface in human skin samples. The tissues were chemically fixed or physically immobilized using <u>h</u>igh-<u>p</u>ressure freezing (HPF) which preserves membranes in their native state (Studer et al., 2008), processed for ultrathin (60 nm) sectioning and analyzed by 2D conventional transmission <u>e</u>lectron <u>m</u>icroscopy (TEM) (**Figures 1E** and **S1D**). The melanocyte-keratinocyte interface revealed numerous plasma membrane-associated cup-shaped invaginations, with a diameter between 43 and 102 nm and an average size of 63.9 nm, that lacked an electron dense cytoplasmic coat (**Figures 1E** and **S1D**, arrowheads). Immunogold labelling on ultrathin cryosections of human skin samples revealed that these invaginations were positive for Cav1 in melanocytes (**Figure S1E**) and were thus identified as caveolae. To access caveolae 3D ultrastructure, thick-sectioned (300 nm) human skin samples were subjected to double-tilt electron tomography (**Figures 1F** and **S1F**). The reconstructed 3D model (**Figure 1F** and **Video 1**) depicts an epidermal area consisting of a transversal section of a melanocyte dendrite (plasma membrane in green) containing pigmented melanosomes (red) and surrounded by a keratinocyte (plasma membrane in blue, presenting keratin bundles on the cytosol). Caveolae (white) were observed in the melanocyte as single or clustered structures known as rosettes (arrowhead and arrow, respectively) that were connected to the cell surface (Richter et al., 2008; Stan, 2005).

3D <u>h</u>uman reconstructed <u>p</u>igmented <u>e</u>pidermis (3D-HRPE) composed of normal human epidermal melanocytes (Mel) and keratinocytes (Ker) are used to study epidermis stratification and pigmentation (Ali et al., 2015). The development of the synthetic tissue includes the initial epidermis stratification at day 4, pigmentation at day 6 and formation of a fully stratified and pigmented epidermis at day 12. To address the distribution and modulation of caveolae during human epidermis formation at cell-cell interface, representative samples of each day were chemically fixed, thin-sectioned and analyzed by conventional TEM (**Figures 1G, H** and **S1G, H**). From day 4 to 12, the melanocyte-keratinocyte interface showed increased numbers of caveolae per 10 µm-length of plasma membrane when compared to homologous keratinocyte-keratinocyte interface (**Figures 1G** and **S1G**). Although the number of caveolae was constant at the melanocyte-keratinocyte interface (**Figure 1G**), differences in caveolae enrichment appeared with time for each cell type (**Figure 1H**). At day 4, when the tissue stratified, caveolae were 4-fold enriched in keratinocytes when compared to melanocytes. However, from day 4 to 6, when the tissue started to pigment, caveolae biogenesis showed a 5-fold increase in melanocytes (**Figure 1H**). As a control, we observed that the number of CCPs, identified by the presence of a characteristic electron dense coat (Heuser, 1980), was similar at both interfaces and cell types and constant over time (**Figures S1H,** bottom panel). This demonstrates that, among these two specialized plasma membrane domains, the melanocyte-keratinocyte interface is preferentially enriched in caveolae. More importantly, during epidermis formation, caveolae numbers are constant at the melanocyte-keratinocyte interface yet they specifically increase in melanocytes when the epidermis starts to pigment suggesting that caveolae could participate in tissue pigmentation.

Ultraviolet (UV) radiations potentiate skin pigmentation by stimulating melanocytes to synthesize and transfer the pigment melanin (Maddodi et al., 2012) while modulating the secretion of keratinocytes signaling factors including exosomes (Lo Cicero et al., 2015; Hirobe, 2005, 2011). We thus examined whether daily low doses of UV-B, which mimic physiological solar exposure (Lo Cicero et al., 2015), could modulate the expression levels of Cav1 in melanocytes and keratinocytes (**Figures 1I** and **S1I**). Cav1 protein levels were increased 6-fold in melanocytes after 3 consecutive irradiations (**Figure 1I**) while keratinocytes only slightly up-regulated Cav1 protein levels in comparison to non-exposed cells (**Figure S1I**). Thus, UV-B exerts a positive role in modulating Cav1 expression in the epidermal unit, yet more prominently in melanocytes. Altogether, we show that melanocytes modulate the levels and distribution of caveolae in response to extracellular and physiological stimuli, such as keratinocytes-secreted factors and UVs.

### Caveolin-1 regulates cAMP production in melanocytes

Considering the prominent function of caveolae in intracellular signaling (Lamaze et al., 2017) and the significant impact of both keratinocyte-secreted factors and UV on caveolae distribution and Cav1 levels, respectively, we investigated whether caveolae-mediated signaling could contribute to pigmentation in melanocytes. Melanocytes express different receptors that activate signal transduction pathways increasing pigmentation (D’Mello et al., 2016; Gordon et al., 1989; Hirobe, 2005, 2014). A key signaling molecule in this process is the second messenger cAMP produced by tmACs downstream of GPCR activation (Buscà and Ballotti, 2000). Interestingly, Cav1 and Cav3 can control cAMP production and were suggested to compartmentalize this second messenger (Allen et al., 2009; Calaghan et al., 2008; Wright et al., 2014). We thus investigated whether Cav1 was required for the production of intracellular cAMP following forskolin (FSK) stimulation, a cell-permeable direct activator of tmACs (Litvin et al., 2003; Metzger and Lindner, 1981; Seamon and Daly, 1981). Melanocytes were treated with control siRNA or siRNAs targeting Cav1 (**Figure S2A**), grown without any cAMP-stimulating molecule and stimulated by FSK (**Figures 2A** and **S2B**). Cav1-depleted melanocytes increased the intracellular cAMP dramatically by 7.5-fold upon stimulation while in control cells, the increase in cAMP was only 3.5-fold (**Figure 2A**). The 2-fold gain in the cAMP production observed in the absence of Cav1 suggests that Cav1 and/or caveolae inhibit tmACs activity in melanocytes. Several studies have reported that caveolae could regulate the activity of various signaling molecules, mostly in an inhibitory fashion, through direct binding to the <u>c</u>aveolin-1 <u>s</u>caffolding <u>d</u>omain (CSD; Lu et al., 2018; Weng et al., 2017). Indeed, the catalytic activity of specific tmACs isoforms can be inhibited by a cell-permeable synthetic peptide which mimics the Cav1 CSD (Toya et al., 1998), and herein after referred to as CavTratin. The stimulation with FSK of CavTratin-treated melanocytes resulted in a 30% reduction of cAMP intracellular levels (**Figures 2B** and **S2C**). These results strongly suggest that caveolin-1 reduces the activity of tmACs and the production of cAMP in melanocytes through direct binding to the Cav1-CSD.

**Figure 2.**
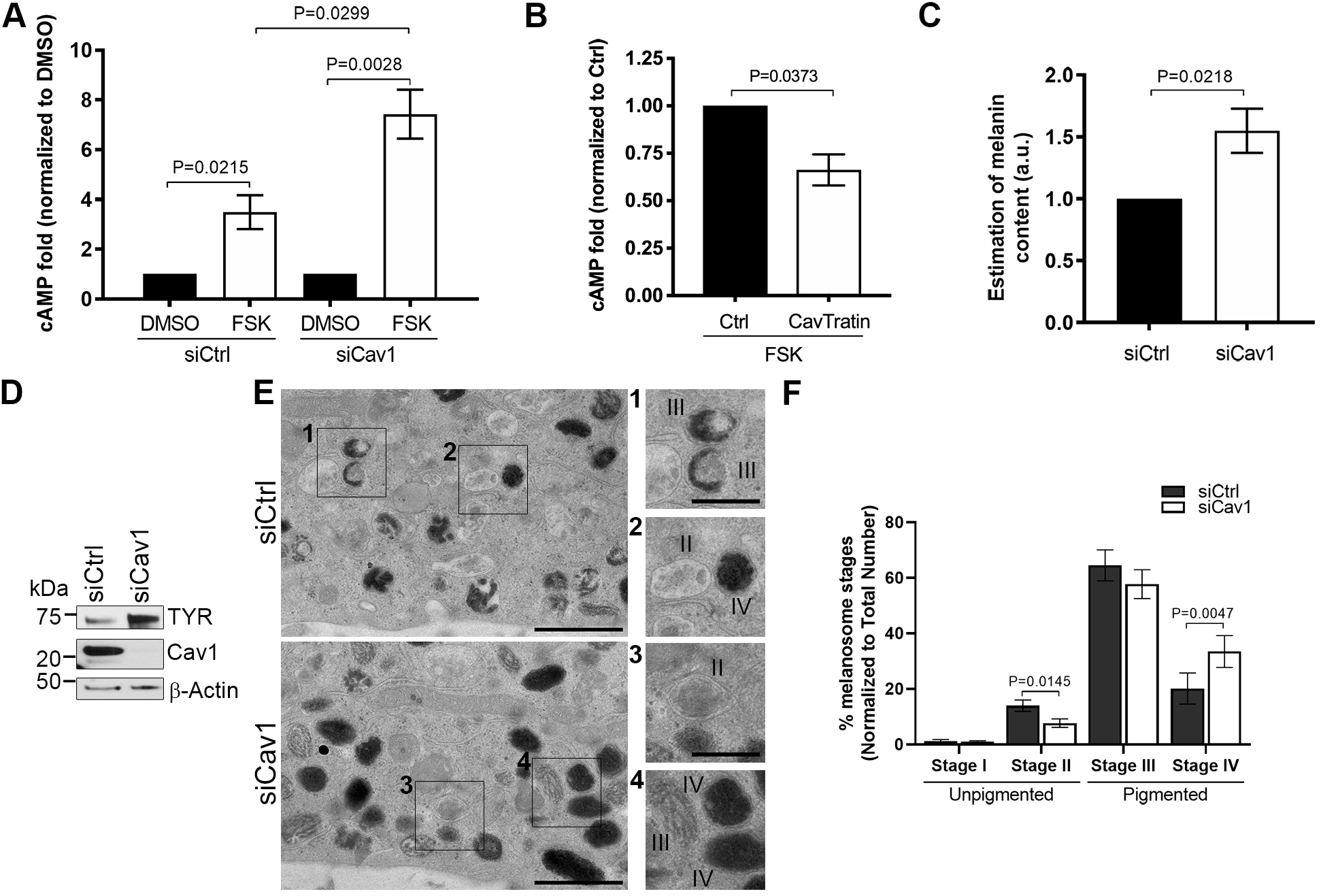
Caveolin-1 depletion in stimulated melanocytes increases cAMP production and pigmentation. **A**. and **B**. Quantification of intracellular cAMP fold-change in melanocytes. A. Melanocytes were transfected with control (Ctrl) or caveolin-1 (Cav1) siRNA for 24h and incubated with DMSO or 30 µM of forskolin (FSK) for 3h (n=3 independent experiments). **B**. Melanocytes were treated with Ctrl (scrambled) or CavTratin (Cav1-scaffolding domain) peptides for 7h and incubated with DMSO or 30 µM of FSK for 1h (Cavtratin: 66.3 ± 8.2; n=3 independent experiments). **C-F**. Melanocytes were treated for 5 days with siCtrl or siCav1. **C**. Estimation of intracellular melanin content (siCav1: 1.5 ± 0.2; n= 4 independent experiments). **D**. Immunoblot analysis of melanocytes lysates probed with the indicated antibodies. ACTB, β-Actin. **E**. Conventional EM images representative of each condition with the respective zooms of the insert regions (left); Bar: original 1 µm, zoomed 0.5 µm; II to IV represent different stages of maturation of melanosomes. **F**. Quantification of the number of non-pigmented (stage I: siCtrl, 1.3 ± 0.5, siCav1, 1.1 ± 0.3; and stage II: siCtrl, 14.0 ± 2.0, siCav1, 7.9 ± 1.6) and pigmented (stage III: siCtrl, 64.5 ± 5.6, siCav1: 58.0 ± 5.2; and stage IV: siCtrl, 20.1 ± 5.6, siCav1, 33.0 ± 5.8) melanosome stages from EM images as in E (n=14 cells each, 4 independent experiments). Values are mean ± s.e.m. A and B, one-way ANOVA with Sidak’s multiple comparison test.

### Caveolin-1 controls pigmentation in melanocytes

In melanocytes, cAMP production by tmACs increases the expression of melanin-synthesizing enzymes that results in increased melanin synthesis (Buscà and Ballotti, 2000; Newton et al., 2007; Pawelek et al., 1973). Growth of melanocytes in supplemented medium containing factors known to elicit intracellular cAMP production (Abdel-Malek et al., 1995; Imokawa et al., 1996), led to a 1.5-fold increase in the intracellular melanin content after Cav1 depletion (**Figures 2C, D** and **S2D**). Melanin synthesis requires the activity of melanogenic enzymes of the tyrosinase family which include the rate-limiting enzyme Tyrosinase (TYR) and the <u>D</u>opa<u>c</u>hrome tautomerase (DCT; Ebanks et al., 2009). In agreement, Cav1-depleted cells showed an enrichment in both TYR and DCT protein levels (**Figures 2D** and **S2E, F**). Within the melanosome, synthesized melanin deposits onto a fibrillar matrix formed upon proteolytic cleavage of the structural protein PMEL (Theos et al., 2005) which expression level remained unchanged in Cav1-depleted melanocytes (**Figure S2G**). Similarly, the expression of the Rab27a GTPase, which regulates melanosome transport to the cell periphery (Bahadoran et al., 2001), was constant (**Figure S2H**). These data indicate that Cav1 depletion specifically affects pigment production in melanosomes, but not their structure nor their intracellular peripheral localization, as also evidenced by conventional TEM of siCav1-treated melanocytes (**Figure S2I**). As pigment production is accompanied by melanosome maturation (Raposo et al., 2001), we used TEM to quantify the early unpigmented (stages I and II) and the mature pigmented melanosomes (stages III and IV) in control and Cav1-depleted melanocytes. Consistent with the biochemical analyses (Figures 2C and D), the number of pigmented stage IV increased significantly with a concomitant decrease in unpigmented stage II in Cav1-depleted melanocytes (Figures 2E and F). Altogether, the caveolin-1 control of early signaling events in melanocytes leads to the regulation of melanin synthesis and melanosome maturation.

### Melanocytes mechanical response to increased cAMP, cell-cell contacts and mechanical stress is regulated by caveolae

Local production of cAMP at the plasma membrane regulates neuronal cell shape (Neves-Zaph, 2017) and epithelial cell polarity (Wojtal et al., 2008). In melanocytes and melanoma cells, the increase of cAMP levels supports dendrite outgrowth (Buscà et al., 1998; Nakazawa et al., 1993; Scott and Leopardi, 2003). For the last few years, caveolae mechanosensing and mechanoprotective functions have emerged as a new major features of caveolae in many cell types *in vitro* and *in vivo* (Sinha et al., 2011; Cheng et al., 2015). In this context, caveolae were recently shown to couple mechanosensing with mechanosignaling in human myotubes (Dewulf et al., 2019). Because Cav1 regulates cAMP levels in melanocytes, we explored the role of caveolae in the mechanical behavior of melanocytes in response to chemical stimulation. Cav1-depleted melanocytes (**Figure S3A**) were grown in three different media: devoid of stimulating molecules (poor medium), containing forskolin (poor medium + FSK) or supplemented with different growth factors (supplemented medium; see Experimental procedures). The shape of the cells was analyzed using fluorescently-labelled phalloidin that stained actin filaments (**Figure 3A**). In the absence of signaling molecules (poor medium), control and Cav1-depleted melanocytes preferentially displayed a similar morphology characterized by the presence of at most two protrusions (Figures 3A and B). Chemical stimulation of control melanocytes increased the number of protrusions, while the majority of Cav1-depleted cells did not extend more than two protrusions (Figures 3A and B). We then characterized the cell morphology by measuring the cell area, major and minor axis and by calculating the length-to-width ratio (**Figures 3C** and **S3B**-**D**). Without chemical stimulation, the length-to-width ratio was similar in control- and Cav1-depleted melanocytes. After stimulation, the area of the cell and the minor axis, but not the major axis, increased in control cells (**Figures S3B-D**). This caused a slight decrease in the length-to width ratio (**Figure 3C**), which reflects cell spreading and formation of dendrite-like protrusions. On the contrary, Cav1-depleted cells responded to stimulation by preserving the cell area (**Figure S3C**) which confirms their elongated shape. Moreover, the major axis increased while the minor axis increased (**Figures S3C** and **D**). This increased dramatically the length-to-width ratio in Cav1-depleted melanocytes (**Figure 3D**) and suggests that cell spreading is mainly occurring along the major axis. Therefore, the sole elevation of intracellular cAMP in melanocytes devoid of caveolae is not sufficient to support the outgrowth of protrusions. Overall, these data indicate that caveolae are required for the mechanical response mediating the morphologic changes of melanocytes to extracellular chemical stimuli.

**Figure 3.**
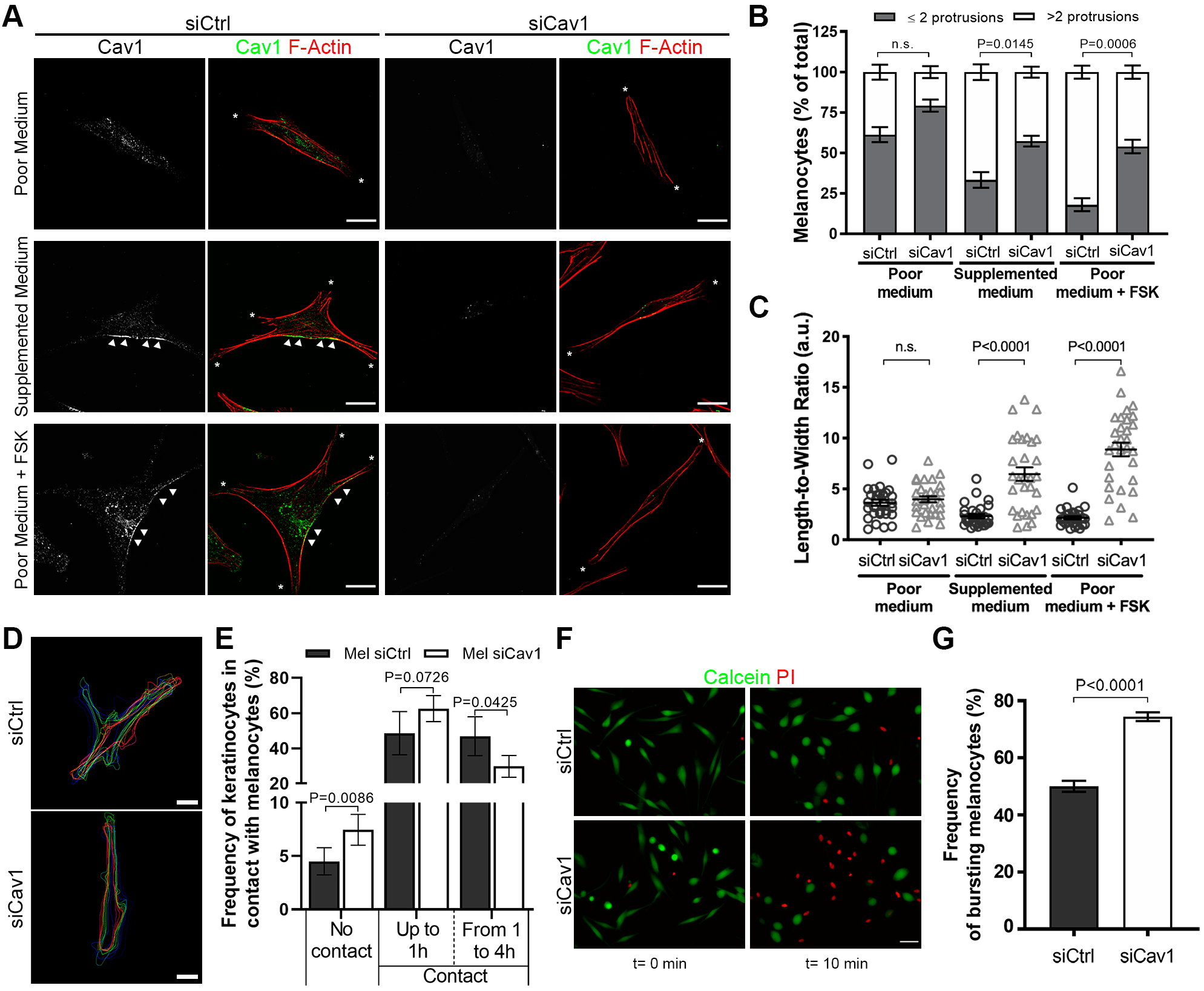
Caveolae contributes to changes in melanocyte morphology, contacts with keratinocytes and mechanoprotection. **A**. IFM images of siCtrl- and siCav1-treated melanocytes incubated with poor medium (+ DMSO), supplemented medium (+ DMSO) or poor medium + 30 µM of FSK for 14h, fixed, immunolabelled for Cav1 (green) and stained for F-actin (phalloidin, red). Arrowheads point Cav1 polarization. Asterisks indicate cell protrusions. Bars, 20 µm. **B**. Frequency of melanocytes showing at most two (≤2) or more than two (>2) membrane protrusions (n=150 cells, 3 independent experiments). **C**. Quantification of the width-to-length ratio of melanocytes cultured as in A (siCtrl: Poor medium, 3.6 ± 0.3, Supplemented medium, 2.3 ± 0.2, Poor medium + FSK, 2.2 ± 0.1; siCav1: Poor medium, 4.0 ± 0.3, Supplemented medium, 6.5 ± 0.7, Poor medium + FSK, 8.9 ± 0.7; n=30 cells, 3 independent experiments). **D** and **E**. Melanocytes treated for 72h with siCtrl or siCav1 were co-cultured with keratinocytes for 14h prior to cell imaging. **D**. Representative projection of time-lapse images with interpolated region of interest for the cell’s boundaries every 20 minutes. Bars, 10 µm. See also **Videos 2** and **3**. **E**. Frequency of keratinocytes contacting melanocytes for a total of 4h (no contact: siCtrl: 4.5 ± 1.3, siCav1: 7.5 ± 1.4; up to 1h: siCtrl: 53.1 ± 11.1, siCav1: 70.2 ± 6.2; from 1 to 4h: siCtrl: 44.1 ± 9.6, siCav1: 26.4 ± 3.9; siCtrl: n=39 videos; siCav1: n=37 videos; 3 independent experiments). **F**. and **G**. Melanocytes treated with siCtrl or siCav1 for 72h were incubated with calcein-AM (green) for 15 minutes, washed and subjected to hypoosmotic shock (30 mOsm) in the presence of propidium iodide (PI, red) for 10 minutes. PI-positive cells (red nuclei) indicate melanocytes with ruptured plasma membrane. See also **Videos 6** and **7**. **F**. First (0 min) and last (10 minutes) still images from the time-lapse acquisition. Bars, 50 µm. **G**. Frequency of bursting melanocytes (siCtrl: 50.0 ± 2.0, n=714; siCav1: 74.4 ± 1.5, n=958; 3 independent experiments). Values are the mean ± s.e.m.

In skin epidermis, the extension of dendrites by melanocytes is crucial to establish contacts with a large number of keratinocytes. To test if caveolae are involved in the change of morphology of melanocytes that occur in response to keratinocytes-secreted factors, we performed time-lapse microscopy of melanocytes co-cultured with keratinocytes. In the absence of direct cell contact with keratinocytes, control melanocytes responded dynamically by extending and retracting dendrite-like protrusions along time (**Video 2**). On the contrary, Cav1-depleted melanocytes displayed an elongated shape and formed fewer projections (**Video 3**). The difference of response due to the absence of caveolae was better evidenced by delineating the cell boundaries during the 4h acquisition (**Figure 3D**) and consistent with the immunofluorescence microscopy data obtained for stimulated melanocytes in monoculture (**Figure 3A**). Besides the established role of extracellular signaling molecules, direct contact between melanocytes and keratinocytes might also promote dendrite outgrowth (Kippenberger et al., 1998). So, we tested if caveolae could contribute to changes in the morphology of the melanocytes in response to cell-cell interactions with keratinocytes. Control melanocytes responded by extending and retracting dendrite-like protrusions when keratinocytes established close contacts (**Video 4**), while Cav1-depleted melanocytes were mostly unresponsive to the contacts made by keratinocytes, formed fewer projections and displayed an elongated shape (**Video 5**). Interestingly, Cav1-depleted melanocytes were more frequently deprived of physical contact by keratinocytes during the total time of acquisition (**Figure 3E**). In contrast, the frequency of melanocytes-keratinocytes contacts that were long-lasting (1-4h) decreased (**Figure 3E** and **Videos 4** and **5**). Thus, melanocytes devoid of caveolae are unable to promote the outgrowth of protrusions in response to the keratinocytes-secreted factors or to the direct contact with keratinocytes. Altogether, this data shows that caveolae in melanocytes play a key role in melanocyte dendrite outgrowth and the establishment and maintenance of contacts with keratinocytes.

The cell mechanical response to changes in shape is correlated with adjustments in the plasma membrane tension to the cytoskeletal architecture and dynamics (Diz-Muñoz et al., 2013; Keren, 2011; Pontes et al., 2017). Under mechanical stress, caveolae serve as a membrane reservoir by disassembling rapidly to buffer variations of plasma membrane tension (Sinha et al., 2011). To address whether the mechanical function of caveolae is involved during the changes in morphology, and thus membrane tension variations, we monitored the resistance of the plasma membrane of melanocytes during membrane tension increase induced by hypoosmotic shock. Melanocytes were pre-incubated with the membrane permeant cytoplasmic green-fluorescent dye calcein-AM and exposed to a 30 mOsm hypo-osmotic shock in the presence of propidium iodide (PI), a non-permeant red-fluorescent DNA intercalating agent. A loss of plasma membrane integrity is revealed by a decrease or absence of the calcein-AM signal whilst acquiring a positive signal for propidium iodide. After 10 min of hypo-osmotic shock, Cav1-depleted melanocytes had burst more frequently than control cells (**Videos 6** and **7** and **Figures 3F** and **3G**), confirming that caveolae offer mechanoprotection to melanocytes experiencing membrane tension variations. All in all, this data indicates that caveolae regulates the mechanical responses of melanocytes observed during contact with keratinocytes or chemical stimuli.

### Loss of caveolae impairs melanin transfer in 2D co-culture and 3D-epidermis

Skin pigmentation relies on the synthesis of the pigment melanin within melanocytes and its transfer to neighboring keratinocytes. Different mechanisms have been proposed for melanin transfer to occur (Tadokoro and Takahashi, 2017; Wu and Hammer, 2014) and all requires the local remodeling of the plasma membrane of melanocytes at the near vicinity of keratinocytes. To address the role of caveolae in melanin transfer, siCtrl- and siCav1-treated melanocytes were co-cultured with keratinocytes for 3 days, after which the cells were analyzed by immunofluorescence (**Figure 4A**). Keratinocytes co-cultured with Cav1-depleted melanocytes were less frequently positive for melanin (**Figure 4B**) and, when positive, showed decreased staining for the pigment (**Figure 4C**). This result shows that caveolae are required for the efficient transfer of melanin from melanocytes to keratinocytes in co-culture.

**Figure 4.**
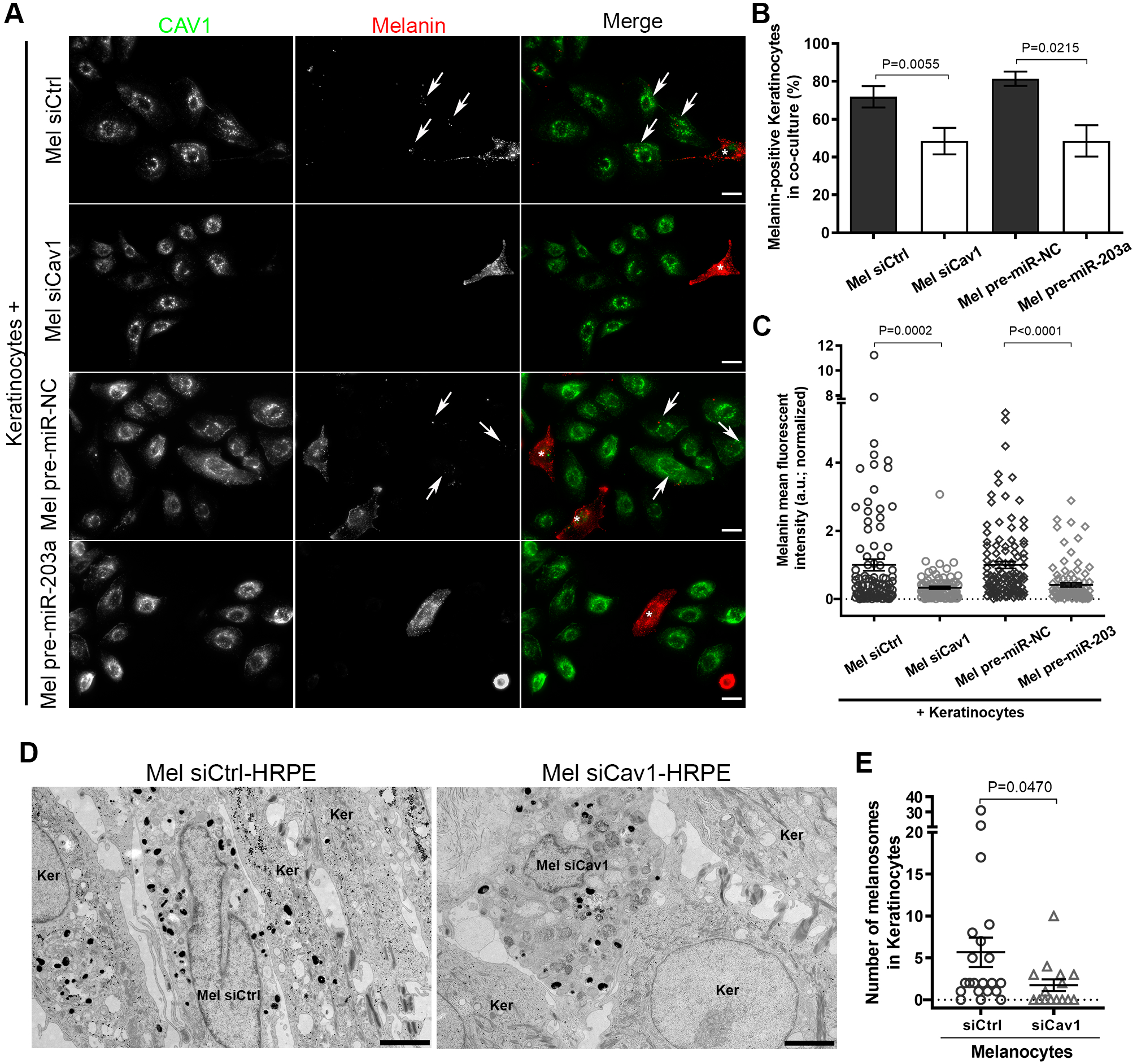
Caveolae in melanocytes are necessary for melanin transfer *in vitro* and in tissue. **A**, **B** and **C**. Melanocytes treated with siCtrl, siCav1, pre-miR-NC (negative control) or pre-miR-203a for 5 days were co-cultured with keratinocytes for the last 2 days. **A**. IFM images of the co-culture immunolabelled for Cav1 (green) and melanin (HMB45, red). Arrows point plasma keratinocytes positive for transferred melanin. Asterisks in merge panels identify melanocytes. Bars, 20 µm. **B**. Quantification of the frequency of keratinocytes positive for melanin in each condition (siCtrl: 71.9 ± 5.7; siCav1: 48.5 ± 7.0; pre-miR-NC: 81.5 ± 3.7; pre- miR-203a: 48.6 ± 8.3; n=150 cells, 3 independent experiments). **C**. Quantification of melanin fluorescent intensity in individual keratinocytes positive for melanin (siCtrl, n=98; siCav1, n=93; pre-miR-NC=111; pre-miR-203a=93; 3 independent experiments). **D**. Conventional EM micrographs of 9 days 3D-HRPE composed of keratinocytes and siCtrl- or siCav1-treated melanocytes. Bars, 2 µm. **E**. Quantification of the number of melanosomes in keratinocytes at the vicinity of melanocytes (siCtrl: 5.7 ± 1.8, n=21 cells; siCav1: 1.7 ± 0.7, n=15 cells; 1 experiment). Values are the mean ± s.e.m.

Interestingly, in melanoma cells, the microRNA-203a (miR-203a) downregulates Cav1 expression (Conde-Perez et al., 2015). Likewise, melanocytes transfected with the pre-mir-203a showed decreased Cav1 protein expression levels (**Figure S4A**). When co-cultured with melanocytes treated with pre-miR-203a, melanin transfer had occurred in fewer keratinocytes (Figures 4A and B), which also showed a decrease content of melanin (**Figure 4C**). The miR-203a is secreted by keratinocytes together with exosomes (Lo Cicero et al., 2015), which suggests that keratinocytes could regulate Cav1 expression levels and caveolae biogenesis in melanocytes to control their signaling and mechanical responses.

Finally, we sought to establish the importance of caveolae in pigment transfer *in vivo*. We turned to the model of skin epidermis (3D-HRPE) and generated three different epidermis composed of keratinocytes either alone (Ker-HRPE) or associated with control or Cav1-depleted melanocytes. The expression of Cav1 mRNAs was efficiently down-regulated after siCav1 treatment in melanocytes (**Figure S4B**). Macroscopic examination of the reconstructed tissue showed unpigmented epidermis when composed of only keratinocytes, and homogenous pigmented epidermis when control melanocytes were added (**Figure S4C**). In contrast, a non-homogenous pigmentation was observed in the epidermis reconstructed with siCav1-treated melanocytes (**Figure S4C**, arrow). The pigmentation defect was further characterized at the ultrastructural level (**Figure 4D**) and revealed that keratinocytes juxtaposed to Cav1-depleted melanocytes contained less melanin than when adjacent to control cells (**Figure 4E**). This data shows that caveolae is a novel player in melanin transfer from melanocytes to keratinocytes in the human epidermis.

## Discussion

Human epidermis pigmentation represents a natural body photo-protective screen that relies on melanocytes and keratinocytes. To adapt to their environment, like during intense solar exposure, these epidermal cells communicate to orchestrate cellular responses important for producing and disseminating the pigment through the tissue. In this study, we provide evidence for a novel physiological role of caveolae in human epidermis pigmentation. By exploiting the signaling and mechanical functions of caveolae, melanocytes respond to the extracellular signals sent by keratinocytes to potentiate skin photo-protection. The capacity of caveolae to modulate intracellular signals, to provide mechano-protection and to support the morphological changes in melanocytes define them as a novel molecular platform required for human skin pigmentation.

Caveolae polarization or enrichment in melanocytes are positively-regulated during the formation of skin, by keratinocytes-secreted factors and by solar mimicking UV-B radiation. Intriguingly, the miR203a secreted together with keratinocytes extracellular vesicles (Lo Cicero et al., 2015) can target Cav1 in melanoma cells (Conde-Perez et al., 2015) and in normal melanocytes. This indicates that keratinocytes directly contribute to fine-tune Cav1 and caveolae in melanocytes so that it cellular responses can be highly organized and coordinated. A down-regulation of Cav1/caveolae would promote pigment production in melanocytes whereas an up-regulation would favor changes in cell morphology and cell-cell contacts, both leading to melanin transfer and skin pigmentation.

Melanocytes devoid of caveolae have higher production of intracellular cAMP after stimulation, whereas treatment with the Cav1 scaffolding domain (CSD) mimicking peptide, CavTratin, has an opposite effect. A classical view of caveolae function in signaling is associated to the intracellular compartmentalization and concentration of different signaling transduction pathways components (Lamaze et al., 2017). In this context, caveolin-1 was shown to regulate the activity of some isoforms of tmACs in cells (Gu et al., 2002; Ostrom et al., 2002). The use of the CavTratin peptide *in vitro* negatively regulated these enzymes with concomitant decrease of cAMP production after stimulation (Toya et al., 1998). This shows that caveolae mitigate the cAMP-dependent signaling in melanocytes, likely through Cav1 binding to tmACs and direct inhibition of their catalytic activity.

In response to increased cAMP production, Cav1-depleted melanocytes do not extend dendritic-like protrusions, strongly suggesting that caveolae couple cAMP-induced signaling to the cell mechanical response. This feature of caveolae might not be only restricted to melanocytes and is likely shared by neural crest-derived cells. Indeed, the modulation of cAMP levels in the vicinity of membrane lipid rafts controls dendritic arborization in mice neurons (Averaimo et al., 2016; Guirland and Zheng, 2007) while neuron-targeted Cav1 enhances branching out of the dendrites (Head et al., 2011; Mandyam et al., 2017). Dendrite outgrowth in human melanocytes and murine melanoma cells is also dependent on cAMP (Buscà et al., 1998; Scott and Leopardi, 2003). Endogenous Cav1 and Cavin1, and therefore caveolae, distribute asymmetrically and cell-autonomously in cultured human melanocytes. Polarization of Cav1 and caveolae is observed in different cells during cell migration (Grande-García and del Pozo, 2008; Navarro et al., 2004). However, cultured melanocytes display a poorly motile behavior, as shown here by time-lapse microscopy, which suggests that caveolae polarization in these cells should perform functions unrelated to cell migration. Melanocytes are likely polarized cells as their shape consists of a cell body facing the basal membrane with multiple dendrites extending upwards and as they express proteins specific of epithelial cells (Valencia et al., 2006). Therefore, we propose that caveolae intrinsic asymmetrical distribution imposes a spatial organization of cAMP-dependent pathways and/or downstream targets in melanocytes that contributes to its polarized organization and ensures its cellular functions.

Caveolae are required for two crucial functions in melanocytes: pigment production and transfer. Stimulation of Cav1-depleted melanocytes causes increased cAMP levels, acceleration of pigment production through likely the up-regulation of Tyrosinase and DCT expression levels. Pigment synthesis and packaging into melanosomes rely on intracellular signaling pathways, among which cAMP synthesis by tmACs is of key importance (D’Mello et al., 2016). The activation of the GPCR-triggered cAMP pathway increases Tyrosinase, TYRP1 and DCT protein content through increased cell transcriptional activity (Bertolotto et al., 1996, 1998a, 1998b) or post-translational events (Abdel-Malek et al., 1995; Newton et al., 2007). This indicates that caveolae key regulation in the production of the pigment occurs through the fine control of cAMP production and downstream pathways.

The fate of melanin in the epidermis is to be transferred to keratinocytes where it shields the nucleus against UV radiations. Here, we establish a correlation between caveolae formation and human skin pigmentation. Caveolae accumulate at melanocyte-keratinocyte interface when the epidermis becomes pigmented while impaired caveolae formation in melanocytes, through Cav1 depletion, decreases melanin transfer in co-culture and reconstructed epidermis. The dendrites of melanocytes are seen as conduits for melanin transfer and points of contact with keratinocytes and, therefore, their plasticity seems important to support these functions. Our results show that caveolae protects the plasma membrane of melanocytes against acute rupture after a mechanical stress thus helping the cells to adjust to tension variations. Several studies illustrate that plasma membrane tension regulates membrane deformations during exo- and endocytosis or changes in cell shape (Dai et al., 1997; Gauthier et al., 2011; Houk et al., 2012; Raucher and Sheetz, 2000). Thus, the dynamic cycle of caveolae mechanics, i.e. disassembly and reassembly, in response to tension variations that occur during melanocytes morphological changes could facilitate both dendrite outgrowth and pigment transfer. Nonetheless, the formation of caveolae and non-caveolae Cav1 clusters could also exert a spatiotemporal control of melanin secretion by favoring the local remodeling of the plasma membrane in response to signaling cues. Therefore, the coupling of signaling and mechanical outputs by caveolae in melanocytes is key to the pigment transfer regulation.

Dysregulation of Cav1 expression in the human skin is associated with hyperproliferative diseases such as melanoma and non-melanoma cancers but also psoriasis (Carè et al., 2011; Gheida et al., 2018; Kruglikov and Scherer, 2019). In melanoma, Cav1 function remains very controversial, since it is recognized as a tumor suppressor and an oncogene (Felicetti et al., 2009; Trimmer et al., 2010). Such discrepancy might be explained by the variations of Cav1 expression during disease progression, as the balance between caveolae signaling and mechanical functions in response to the extracellular environment changes during tumor mass growth (Lin et al., 2007). Long-term exposure to UV radiations is a key factor causing skin cancers (MacKie, 2006) and high levels of expression of the miR-203a occurs in psoriatic lesions (Huang et al., 2015). We, thus propose caveolae as a novel modulator of skin pigmentation that couple signaling with mechanical responses in melanocytes. The characterization of the physiology underlying these two caveolae functions, by and in response to the extracellular context, will enable to decipher its defects and associated consequences in disease.

## Acknowledgments

We thank the Structure and Membrane Compartment laboratory for insightful discussions, Lucie Sengmanivong from the Nikon Imaging Centre at Institut Curie-CNRS for help in image acquisition, and Gisela D’Angelo (Institut Curie, Paris, France) and Corinne Bertolotto (Université Côte d’Azur, Nice, France) for critical reading of the manuscript. The authors acknowledge the Nikon Imaging Center at Institut Curie/ Centre National de la Recherche Scientifique and the PICT-IBiSA, a member of the France-BioImaging national research infrastructure (ANR10-INBS-04). This work has received support under the program “Investissement d’Avenir” launched by the French Government and implemented by the Agence Nationale de la Recherche (ANR) with the references ANR-10-LBX-0038 and ANR-10-IDEX-0001-02 PSL, Fondation pour la Recherche Médicale (Equipe FRM DEQ20140329491 Team label to G.R.), Agence Nationale de la Recherche (“MOTICAV” ANR-17-CE13-0020-01 to C.L.), the Fondation ARC pour la Recherche sur le Cancer (PJA20161204965 to C.D., and Programme Labellisé PGA1-RF20170205456 to C.L.), Labex CelTisPhyBio (to L.D.), Groupe Clarins, Institut Curie, CNRS and INSERM. The authors declare no competing financial interests.

## Author contribution

Conceptualization, L.D., C.B., C.G., K. V., G.R., C. L. and C.D.; Methodology, L.D., I.H., N.A., M.D. and C.D.; Formal analysis, L.D.; Investigation, L.D., I.H., F.G-M., N.A., M.D., M.R., C.V.L. and C.D.; Visualization, L.D.; Writing – Original Draft, L.D. and C.D.; Writing – Review and Editing, All authors; Project Administration and Funding Acquisition, G.R., C.L. and C.D.; Supervision, L.D., C.B., C.G., K.V., G.R., C.L. and C.D.

## Experimental procedures

### Antibodies

The following antibodies were used for immunoblot (IB) or immunofluorescence (IFM): rabbit polyclonal anti-Caveolin1 (BD Transduction Laboratories; 1:5000 [IB]; 1:200 [IFM]); rabbit polyclonal anti-PTRF (CAVIN-1; Abcam; 1:200 [IFM]); mouse monoclonal anti-HMB45 (recognizing PMEL-positive fibrils onto which melanin deposits, here used as a melanin marker; clone HMB45; abcam; 1:200 [IFM]); mouse monoclonal anti-α adaptin (clone AP6; abcam; 1:50 [IFM]); sheep polyclonal anti-EGFR (Fitzgerald; 1:400 [IFM]); mouse monoclonal anti-Tyrosinase (clone T311; Santa Cruz biotechnology; 1:200 [IB]); mouse monoclonal anti-DCT (clone C-9; Santa Cruz biotechnology; 1:200 [IB]); rabbit polyclonal anti-Pep13h (Raposo et al., 2001; 1:200 [IB]); goat polyclonal anti-Rab27a (SICGEN; 1:1000 [IB]); mouse monoclonal anti-ACTB (β-actin; clone AC-74; Sigma; 1:2000; [IB]); rabbit polyclonal anti-GAPDH (Sigma; 1:10000; [IB]); rabbit polyclonal anti-Calnexin (Enzo Life Sciences; 1:1000; [IB]). Secondary antibodies coupled to horseradish peroxidase (HRP) were used at 1:10000 for IB (Abcam). Secondary antibodies and phalloidin conjugated to 488, 555 and 647 Alexa dyes were used at 1:200 (Invitrogen) for IFM.

### Cell culture

#### Primary cells

Normal Human Epidermal Melanocytes and Normal Human Epidermal Keratinocytes used in this study were isolated from neonatal foreskins and purchased from CellSystems, Sterlab and PromoCell. Melanocytes and keratinocytes were used from passage two and five and maintained in culture in DermaLife Basal Medium supplemented with DermaLife M Life factors (Melanocytes supplemented medium) or in DermaLife Basal Medium supplemented with DermaLife K Life, respectively.

#### Cell line

HeLa cells were cultured in DMEM supplemented with 10% (v/v) FBS, 100 U/ml penicillin G and 100 mg/ml streptomycin sulfate (Gibco). All cells were maintained at 37°C in a 5% (v/v) CO_2_ incubator.

### siRNA and miRNA transfections

For melanocytes siRNA and miRNA transfections, cells were seeded in the appropriate wells or plates and transfected with 0,2 µM of siRNA using Oligofectamine (Invitrogen) accordingly to manufacturer’s instructions using non-targeting siRNA (siCtrl; 5’-AATTCTCCGAACGTGTCACGT-3’) and siRNA targeting Cav1 (SI00299635 and SI00299628) from Qiagen, or using pre-miR-NC (negative control; #AM17111) and pre-miR-203a (#AM17100) from Thermo Fischer Scientific. In 3D-HRPE experiments, melanocytes were transfected previously to reconstruction with 1 µM of siRNA using DharmaFECT and following the manufacturer’s protocol (Dharmacon, Horizon) using non-targeting siRNA (Accell non-targeting pool) or siRNA targeting Cav1 (SMARTpool: Accell Cav1) from Dharmacon.

### Co-cultures and media incubation

#### Co-cultures

Melanocytes and keratinocytes or HeLa were seeded in the following ratio, respectively: 1:4 for 24h before fixation to quantify caveolae polarization (Figure 1); at 1:4 for 14h before time-lapse acquisition (Figure 3); and 1:1 for 3 days before fixation to quantify melanin transfer (Figure 4). All co-cultures were done in Melanocytes supplemented medium.

#### Media incubation

Keratinocytes medium from a confluent flask in culture for 48h was collected and centrifuged at 200 rcf to remove cell debris. The Keratinocytes-conditioned medium (Ker-CM) was immediately used or stored at −80°C (Figure 1). Melanocytes were seeded in melanocytes supplemented medium for 6h after which, this medium was removed, the cells washed in phosphate-buffered saline (PBS) and poor medium, poor medium supplemented with 30 µM of forskolin (FSK, Sigma), new melanocytes supplemented medium or Ker-CM was added and kept for approximately 14h before fixation. Dimethylsulfoxide (DMSO) was added to the medium as a control to FSK addition.

### UV treatment

Melanocytes and Keratinocytes were seeded in six-well plates at day 0 and irradiated with a single shot of 10 mJcm-2 of ultraviolet B (312 nm) during 3 consecutive days using a Biosun machine (Vilber Lourmat, Suarlée, Belgium). Cell medium was replaced by PBS before irradiation and replaced by the culture medium just after the treatment. The cells were then incubated overnight and recovered by trypsinization at the indicated time points.

### Skin samples

Healthy skin samples were obtained from surgical left-over residues of breast or abdominal reduction from healthy women. Written informed consent was obtained in accordance with the Helsinski Declaration and with article L.1243-4 of the French Public Health Code. Given its special nature, surgical residue is subject to specific legislation included in the French Code of Public Health (anonymity, gratuity, sanitary/safety rules…). This legislation does not require prior authorization by an ethics committee for sampling or use of surgical waste (http://www.ethique.sorbonne-paris-cite.fr/?q=node/1767).

### Human Reconstructed Epidermis (3D-HRPE)

The following protocol was adapted from (Salducci et al., 2014). Briefly, dead de-epidermized dermis were prepared as follows. Skin samples from healthy adults were obtained, cut in circular pieces (18 mm diameter) and incubated 20 min at 56°C in HBSS (Invitrogen) containing 0,01% (v/v) Penicillin/Streptomycin (Invitrogen). Epidermis was removed and collected dermis fragments were sterilized in 70° Ethanol, washed twice in HBSS, frozen in HBSS (−20°C) and submitted to six cycles of freezing-thawing to eliminate fibroblasts. De-epidermized dermis were then placed at the bottom of a 6-well plate in 3D-HRPE culture medium composed of IMDM medium (Invitrogen) and keratinocytes medium (CellSystems) at a proportion of 2/3 to 1/3, respectively, and containing 10% (v/v) of calf fetal serum gold (PAA). siRNA-treated melanocytes and non-treated keratinocytes were seeded at a proportion 1:20, respectively, in a culture insert of 8 mm of diameter affixed on the dermis to promote cell adhesion. After 24h, the culture insert was removed and the de-epidermized dermis submerged for 3 days in 3D-HRPE culture medium to promote cell proliferation. Tissue stratification was initiated by moving up the de-epidermized dermis to the air-liquid interface. At day 4, the newly formed epidermis started to stratify, at day 6 it started to pigment and at day 9 to 12, the epidermis was fully stratified and pigmented. All the incubation steps were performed at 37°C in a 5% CO_2_ incubator.

### Measurement of intracellular cAMP levels

Melanocytes were transfected once with the indicated siRNAs and cultured in DermaLife Basal Medium without the addition of StiMel8 LifeFactor (Poor medium) for 24h. DMSO or 30 µM of FSK were added to the respective wells for 3h after which the cells were collected and the intracellular cAMP content measured using the cAMP complete ELISA kit (Enzo Life Sciences) following manufacturer’s instructions. For the treatment with the peptides, NHEMs were maintained in Poor medium for 14h before the addition of the peptides Ctrl (scrambled sequence) or CavTratin (Cav1-scaffolding domain, CSD) during 7h. Then the cells were incubated for 1h with DMSO or 30 µM of FSK after which the cells were collected and the intracellular cAMP content measured.

### Melanin assay

Melanocytes were transfected twice at days 1 and 3 for a total of 5 days with the indicated siRNAs. Cells were then collected, sonicated in 50mM Tris-HCl pH 7.4, 2mM EDTA, 150mM NaCl, 1mM dithiothreitol (with the addition of protease inhibitor cocktail, Roche) and pelleted at 20,000g for 15 min at 4°C. The pigment was rinsed once in ethanol:ether (1:1) and dissolved in 2M NaOH with 20% (v/v) DMSO at 60°C. Melanin content was measured by optical density at 490 nm (Spectramax 250, Molecular Devices).

### Membrane bursting assay

Melanocytes were transfected twice with the indicated siRNAs at day 1 and day 3 for a total of 3 days and seeded in 12-well plates for 24h in supplemented medium. At day 4, cells were incubated in 5 µg/ml of Calcein-AM (Life technologies) for 15 min at 37°C protected from light. The wells were washed once with melanocytes supplemented medium and maintained until image acquisition. Melanocytes supplemented medium was diluted in 90% (v/v) water, the equivalent of 30 mOsm hypo-osmotic shock, followed by the addition of 2 mg/mL of propidium iodide (PI, Sigma) and used to induce the plasma membrane (Dewulf et al., 2019). Immediately after medium replacement, images were acquired every minute for a total of 10 min in an inverted microscope (Eclipse Ti-E, Nikon), equipped with a CoolSnap HQ2 camera, using the 20x 0.75 NA Plan Fluor dry objective together with MetaMorph software (MDS Analytical Tecnhologies).

### Melanin transfer assay

The detailed protocol for the melanin transfer assay is described elsewhere (Ripoll et al., 2018). Melanocytes were transfected twice with the indicated siRNA or miRNAs at day 1 and day 3 for a total of 5 days. At day 3, Melanocytes were co-cultured with keratinocytes for a total of 2 days. Images were acquired with an upright epi-fluorescence microscope (Eclipse Ni-E, Nikon) equipped with a CoolSnap HQ2 camera, using a 40x 1.4 NA Plan Apo oil immersion objective together with MetaMorph software.

### Immunofluorescence microscopy

Cell monolayers seeded on glass coverslips were fixed with 4% (v/v) paraformaldehyde in PBS at room temperature for 15 minutes, then washed three times in PBS and once in PBS containing 50 mM glycine. Primary and secondary antibodies dilutions were prepared in the buffer A: PBS containing 0,2% (w/v) BSA and 0,1% (w/v) saponin. The coverslips were washed once in the buffer A and after incubated for 1 h at room temperature (RT) with the primary antibodies. Following one wash step in buffer A, the coverslips were incubated for 30 minutes at RT with the secondary antibodies. If staining with phalloidin was included, the coverslips were washed in buffer A and incubated in the same buffer with phalloidin at 4°C during 14h.The final wash step was done once in the buffer A, once in PBS and once in water. The coverslips were mounted onto glass slides using ProLong™ Gold Antifade Mount with DAPI (ThermoFischer Scientific). Images were acquired on an Applied Precision DeltavisionCORE system (unless stated otherwise), mounted on an Olympus inverted microscope, equipped with a CoolSnap HQ2 camera (Photometrics), using the 40x 1.3 NA UPLFLN or the 60x 1.42 NA PLAPON-PH oil immersion objectives. Images were deconvolved with Applied Precision’s softWorx software (GE Healthcare).

### Time-lapse microscopy

Melanocytes were transfected twice with the indicated siRNA molecules at day 1 and day 3 for a total of 3 days and co-cultured with keratinocytes in an ibidi polymer coverslip µ-slide (Ibidi) for 14h before imaging. Images were acquired every 5 min for a total of 240 min in an inverted microscope (Eclipse Ti-E, Nikon), equipped with a CoolSnap HQ2 camera, using the 40x 0.75 NA Plan Fluor dry objective together with NIS-Elements software (Nikon).

### Electron microscopy

#### Conventional EM

Human skin epidermis tissues and 3D-HRPE were prepared for EM as described here (Hurbain et al., 2018). For high-pressure freezing, the tissue was high-pressure frozen using an HPM 100 (Leica Microsystems) in FBS serving as filler and transferred to an AFS (Leica Microsystems) with precooled (−90°C) anhydrous acetone containing 2% (v/v) osmium tetroxide and 1% (v/v) of water. Freeze substitution and Epon embedding was performed as described in (Hurbain et al., 2008). For chemical fixation, melanocytes seeded on coverslips and transfected twice with the indicated siRNAs at days 1 and 3 for a total of 5 days were fixed in 2.5 % (v/v) glutaraldehyde in 0.1M cacodylate buffer for 24h, post-fixed with 1% (w/v) osmium tetroxide supplemented with 1.5% (w/v) potassium ferrocyanide, dehydrated in ethanol and embedded in Epon as described in (Raposo et al., 2001). Ultrathin sections of cell monolayers or tissue were prepared with a Reichert UltracutS ultramicrotome (Leica Microsystems) and contrasted with uranyl acetate and lead citrate.

#### Electron tomography

300 nm thick sections were randomly labeled on the two sides with 10 nm Protein-A gold (PAG). Tilt series (2 perpendicular series, angular range from −60° to +60° with 1° increment) were acquired with à Tecnai 20 electron microscope (ThermoFischer Scientific). Projection images (2048 x 2048 pixels) were acquired with a TEMCAM F416 4k CMOS camera (TVIPS). Tilt series alignment and tomogram computing (resolution-weighted back projection) were performed using etomo [IMOD –(Mastronarde, 1997)] software. PAG 10 nm at the surface of the sections was used as fiducial markers. Manual contouring of the structures of interest was performed using IMOD (Kremer et al., 1996).

#### Immuno-EM

Cell samples were fixed with 2% PFA in a 0.1M phosphate buffer pH7.4 and processed for ultracryomicrotomy as described (Hurbain et al., 2017). Ultrathin sections were prepared with an ultracryomicrotome UC7 FCS (Leica) and underwent single immunogold labeling with protein A conjugated to gold particles 10 nm in diameter (Cell Microscopy Center, Department of Cell Biology, Utrecht University). All images were acquired with a Transmission Electron Microscope (Tecnai Spirit G2; ThermoFischer Scientific, Eindhoven, The Netherlands) equipped with a 4k CCD camera (Quemesa, EMSIS, Muenster, Germany).

### Image analysis and quantifications

#### Conventional EM

Caveolae and clathrin-coated pits (Stan, 2005), and melanosome stages were identified based on their ultrastructural features (Raposo et al., 2001). Caveolae structures associated with plasma membranes of randomly selected cell profiles were quantified from 2-D ultrathin sections of 3D-HRPE. The length of the plasma membranes either of melanocytes or keratinocytes were measured using ITEM software (EMSIS) and the total number of caveolae found associated was reported to 10 µm of plasma membrane of the respective cell type. For melanosome stage quantification, the areas corresponding to the tips of the cells were not considered.

#### Immunoblot

Quantification of protein content on western blot was performed using Fiji software, the background subtracted and intensities were normalized to loading control.

#### Caveolae asymmetric distribution by IFM

Images of endogenous staining for Cav1 and Cavin1 polarized in co-culture were acquired and the background subtracted. Two identical boxes were positioned at the plasma membrane but on opposite sides of the cells and the average fluorescent intensity retrieved. The frequency of Cav1 and Cavin1 polarization in melanocytes was defined by identifying cells with one side presenting enriched labelling closely associated with the plasma membrane.

#### Protrusions and cell morphology

A protrusion was defined as an actin-stained extension originated from the soma of the cell. Isolated cells-treated with siCtrl and siCav1 were selected randomly, imaged and the size parameters (area, length-to-width ratio, major and minor axis) were retrieved. The contour of the cell was done using the wand tool and corrected manually if needed recurring to the tool OR (combine).

#### Time of contact

A cell-cell contact was defined optically when the plasma membrane of keratinocyte and melanocyte directly contacted, excluding filopodia.

#### Cell boundary in time-lapse microscopy

Melanocytes cell contour was drawn manually every 5 frames and, in between those frames, the tool Interpolate ROI was used. When needed, the cell boundary was adjusted manually.

#### Membrane bursting assay

The background of time-lapse images acquired from the different channels – PI (mcherry) and Calcein-AM (gfp) – was removed with the tool subtract background from Fiji software and cell’s burst determined when the nuclei was red-stained with concomitant loss of gfp at the cytoplasm.

#### Melanin transfer assay

Image analysis and quantifications are described elsewhere (Ripoll et al., 2018). All images are maximum-intensity z projections of three-dimensional image stacks acquired every 0.2 µm. Fiji software was used for image analysis.

### Immunoblot

Cells analyzed by immunoblot were collected by trypsinization followed by centrifugation. The cell pellet was resuspended in lysis buffer (20 mM Tris-HCl pH 7.2, 150 mM NaCl, 0.1% (v/v) Triton X-100) containing a protease inhibitor cocktail (Roche). The protein content of the lysates was determined with the Pierce™ BCA Protein Assay Kit (ThermoFischer Scientific), the concentrations adjusted with loading buffer (250 mM Tris-HCl pH 6.8, 10% (v/v) SDS, 50% (v/v) Glycerol, 0.5 M β-mercaptoethanol, 0.5% (w/v) Bromophenol blue) and the samples boiled for 5 min at 95°C. After SDS-PAGE using NuPage (4-12%) Bis-Tris gels (Invitrogen), the proteins were transferred to 0.2 μm pore-size nitrocellulose membranes (Millipore) and blocked in PBS with 0.1% (v/v) Tween and 4% (w/v) non-fat dried milk. The membranes were then incubated with the indicated primary antibodies prepared following manufacturer’s instructions. The detection was done using HRP-conjugated secondary antibodies, ECL Plus Western blotting detection system (GE Healthcare) and exposure to Amersham Hyperfilm ECL (GE Healthcare).

### Quantitative real-time PCR

Melanocytes transfected once with the indicated siRNAs for a total of 12 days were collected at days 1 and day 12. The RNA was extracted using the Qiagen RNeasy Mini Kit for RNA extraction (Qiagen) and the cDNA generated using the Transcriptor Universal cDNA Master (Roche) following manufacturer’s protocols. 0.3 µg of RNA was used for the quantitative real-time PCR, the mix prepared accordingly to Probes Master (Roche) and the RealTime ready Custom Panels plates (Roche) used for the assay. The method ΔΔCT was used to obtain the relative expression levels and the ratio between the control and gene of interest was calculated with the formula 2-ΔΔCT.

### Statistical analysis

All the statistical analysis on the collected data was performed using GraphPad Prism, version 7 and 8, GraphPad Software, San Diego California, USA (www.graphpad.com). Scored or quantified cells in each experiment were randomly selected, and all experiments were repeated at least three times unless stated otherwise. Results are reported as mean ± standard error of the mean (s.e.m.). Statistical analysis between three or more experimental groups was performed with one-way ANOVA and Tukey’s multiple comparison test while for comparisons between two sets of data it was used the two-tailed unpaired Student’s t-test with Welch’s correction (unless stated otherwise in figure legends). Differences between data sets were considered significant if P < 0.05.

## Supplemental Information

**Figure S1 – Related to Figure 1. A.** IFM images of melanocytes (white asterisks) in mono- or in co-culture with HeLa cells for 1 day, fixed and immunolabelled for Cav1 or Cavin1 (top or bottom, respectively; green) and melanin (HMB45, red). Asterisks represent melanocytes. **B**. IFM images of a melanocyte immunolabelled for Cav1 (green) and AP-2 (red). **C**. IFM images of melanocytes grown in Ker-CM for approximately 14h, fixed and Cav1 or Cavin1 (top or bottom, respectively; green) and melanin (HMB45, red). B and C. Arrowheads point Cav1 and Cavin1 polarization, Bars, 10 µm. **D**. Raw EM micrographs of human skin epidermis chemically fixed as represented in Figure 1A. Arrowheads point plasma membrane invaginations with morphological features of caveolae. The boxed regions mark the area zoomed in the insets in Figure 1A. Bar, 1 µm. **E**. Ultrathin cryosection of human skin epidermis immunogold labelled for Cav1 (PAG10nm). The boxed region marks the area zoomed in the inset. Bars: original, 1 µm; zoom, 250 nm. **F**. Slices of the electron tomographic reconstruction depicting the Mel-Ker interface shown in Figure 1B. Large electron dense (black) structures correspond to melanin and arrows point plasma membrane invaginations with morphological features of caveolae. Bar, 1 µm. See also **Video 1**. **G**. Conventional EM micrographs of 3D-HRPE at day 6 showing keratinocyte-keratinocyte (top) or melanocyte-keratinocyte (bottom) interfaces. Arrowheads point plasma membrane invaginations with morphological features of caveolae. The boxed region marks the area zoomed in the inset below. Bars: original, 1 µm; zoom 0.5 µm. **H**. Quantification of the number of CCP profiles per 10 µm of plasma membrane at the indicated interfaces (top) or cell type at melanocyte-keratinocyte interface (bottom) (top, Mel-Ker: day4, n=28; day6, n=26; day 12, n=20; Ker-Ker: day4, n=12; day6, n=8; day12, n=10; bottom, Melanocytes or Keratinocytes: day4, n=14; day6, n=13; day 12, n=10; n=number of interfaces). **I**. Immunoblot analysis and quantification (n=2 independent experiments) of Cav1 expression levels in keratinocytes exposed to daily radiations of UV-B (10 mJ/cm^2^) in three consecutive days. H and I, data are presented as mean ± s.e.m.

**Figure S2 – Related to Figure 2. A**. Immunoblot analysis of Cav1 expression levels in melanocytes treated with siCtrl or siCav1 for 24h (left) and associated quantification (right; siCav1: 11.3 ± 4.4; n= 3 experiments). **B**. Quantification of cAMP intracellular concentration in melanocytes treated with siCtrl or siCav1 and incubated with DMSO or 30 µM of FSK for 3h (siCtrl + DMSO: 1.6 ± 1.0; siCtrl + FSK: 4.4 ± 1.8; siCav1 + DMSO: 2.0 ± 1.6; siCav1 + FSK: 8.3 ± 4.6; n=3 independent experiments). **C**. Quantification of intracellular cAMP fold-change in melanocytes treated with Ctrl and CavTratin (Cav1 scaffolding domain) peptides for 7h and incubated with DMSO or 30 µM of FSK for 1h (Ctrl + FSK: 3.7 ± 0.4; CavTratin + FSK: 2.5 ± 0.5; n= 3 experiments). **D**. Quantification of TYR and Cav1 protein levels shown in Figure 2D (siCav1, TYR: 167.3 ± 25.4; Cav1: 11.4 ± 1.2). **E-G**. Immunoblot analysis of melanocytes treated for 5 days with siCtrl or siCav1 using the indicated antibodies (left) and associated quantifications (right; siCav1, DCT: 484.3 ± 84.6; PMEL: 90.4 ± 13.6; Rab27a: 115.3 ± 24.4). **H**. Conventional EM images representative of each condition. Bars: 2 µm. Quantifications of protein expression levels were done relative to the loading control and normalized to siCtrl treated cells (n=3 independent experiments). Values are the mean ± s.e.m.

**Figure S3 – Related to Figure 3. A.** Immunoblot analysis of Cav1 expression levels in melanocytes treated 48h with siCtrl or siCav1 (left) and associated quantification (right; siCav1: 2.0 ± 1.2; n=3 independent experiments). **B**., **C**. and **D**. Quantification of the area (B), major axis (C) and minor axis (D) of siCtrl- and siCav1-treated melanocytes grown in the conditions described in figure 3A (n=30 cells, 3 independent experiments). **B**. siCtrl: Poor medium, 998.2 ± 68.4, Supplemented medium, 1644 ± 73.2, Poor medium + FSK, 1501 ± 95.9; siCav1: Poor medium, 944.9 ± 61.0, Supplemented medium, 1092 ± 64.3, Poor medium + FSK, 941.4 ± 63. **C**. siCtrl: Poor medium, 64.6 ± 3.1, Supplemented medium, 67.2 ± 2.5, Poor medium + FSK, 61.6 ± 2.1; siCav1: Poor medium, 66.6 ± 3.3, Supplemented medium, 87.4 ± 4.2, Poor medium + FSK, 97.6 ± 4.5. **D**. siCtrl: Poor medium, 20.2 ± 1.4, Supplemented medium, 32.0 ± 1.6, Poor medium + FSK, 31.1 ± 1.6; siCav1: Poor medium, 18.5 ± 1.0, Supplemented medium, 17.5 ± 1.5, Poor medium + FSK, 12.8 ± 1.0. **E**. Immunoblot analysis of Cav1 expression levels in melanocytes treated 72h with siCtrl and siCav1 (left) and associated quantification (right; siCav1: 9.6 ± 4.3; n=3 independent experiments). Values are the mean ± s.e.m.

**Figure S4 – Related to Figure 4. A.** Immunoblot analysis of Cav1 expression levels in melanocytes treated 5 days with pre-miR-NC or pre-miR-203a (left) and associated quantification (right; pre-miR-203a: 25.9 ± 9.3; n=3 independent experiments). **B**. Cav1 mRNA levels in melanocytes treated 9 days with siCtrl or siCav1 were analyzed by quantitative RT-PCR (siCav1: 0.41). **C**. Macroscopic images of 3D-HRPE reconstructed with keratinocytes alone (left), keratinocytes and siCtrl-treated melanocytes (middle) or keratinocytes and siCav1-treated melanocytes (right). Arrow points towards the de-pigmented area. Values are the mean ± s.e.m.

**Video 1** – Ultrastructural 3D-model of a Melanocyte-Keratinocyte interface at a human skin epidermis by electron tomography; melanocytes plasma membrane (green), keratinocytes plasma membrane (blue), pigmented melanosomes (red), melanin pigment (black) and caveolae (white) in single and clustered structures. See also Figures 1F and S1F.

**Video 2** – Time-lapse microscopy of siCtrl-treated melanocytes co-cultured with keratinocytes used to draw the cell boundaries (see also Figure 3D, top). Trans-illumination. Acquisition parameter: 200 ms exposure. Video is shown at 7 frames/second. Bar, 20 µm.

**Video 3** - Time-lapse microscopy of siCav1-treated melanocytes co-cultured with keratinocytes used to draw cell boundaries (see also Figure 3D, bottom). Trans-illumination. Acquisition parameter: 200 ms exposure. Video is shown at 7 frames/second. Bar, 20 µm.

**Video 4** – Time-lapse microscopy of siCtrl-treated melanocytes (contoured in yellow, left) co-cultured with keratinocytes (contoured in green, left). See also Figure 3E. Trans-illumination. Acquisition parameter: 200 ms exposure. Video is shown at 7 frames/second. Bar, 25 µm.

**Video 5** – Time-lapse microscopy of siCav1-treated melanocytes (contoured in yellow) co-cultured with keratinocytes (contoured in green). See also Figure 3E. Trans-illumination. Acquisition parameter: 200 ms exposure. Video is shown at 7 frames/second. Bar, 25 µm.

**Video 6** – Time-lapse microscopy of the burst assay for siCtrl melanocytes. Acquisition parameters: 80-150 ms. Video is shown at 7 frames/second. Bar, 50µm.

**Video 7** – Time-lapse microscopy of the burst assay for siCav1melanocytes. Acquisition parameters: 80-150 ms. Video is shown at 7 frames/second. Bar, 50µm.

