## Supplementary figures and images for "Caveolae coupling of melanocytes signaling and mechanics is required for human skin pigmentation"

### Supplemental Figure 1

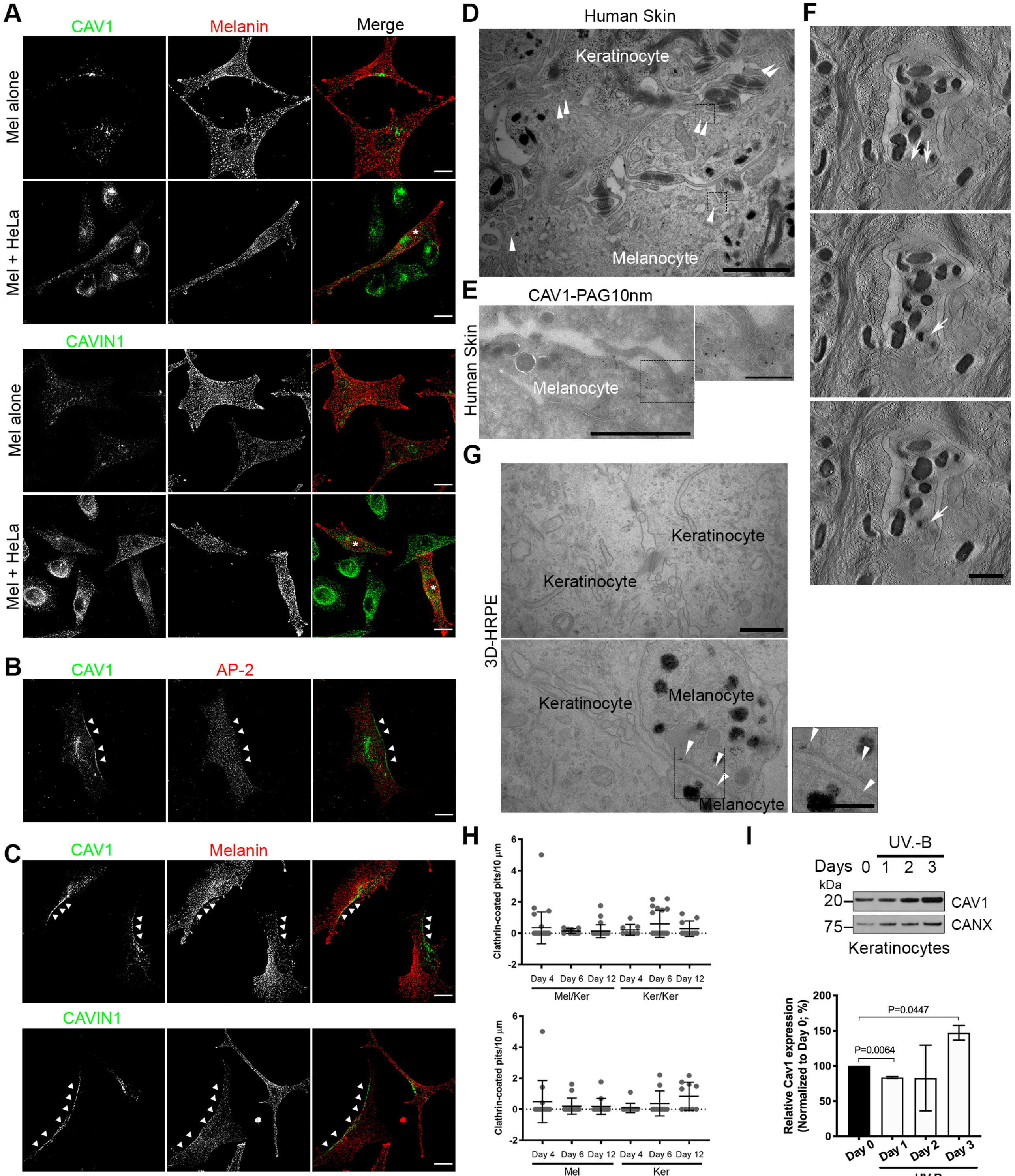

### Supplemental Figure 2

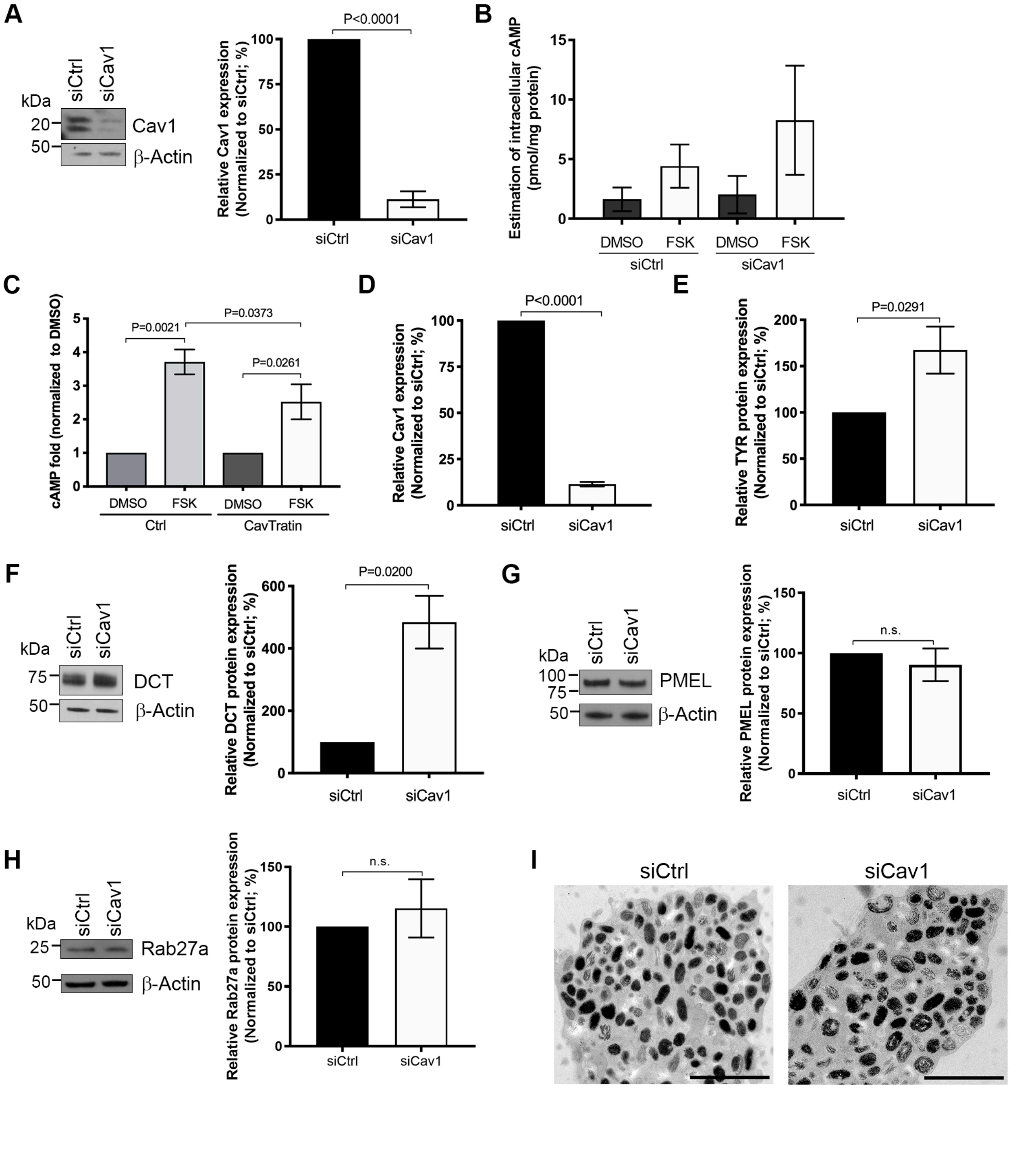

### Supplemental Figure 3

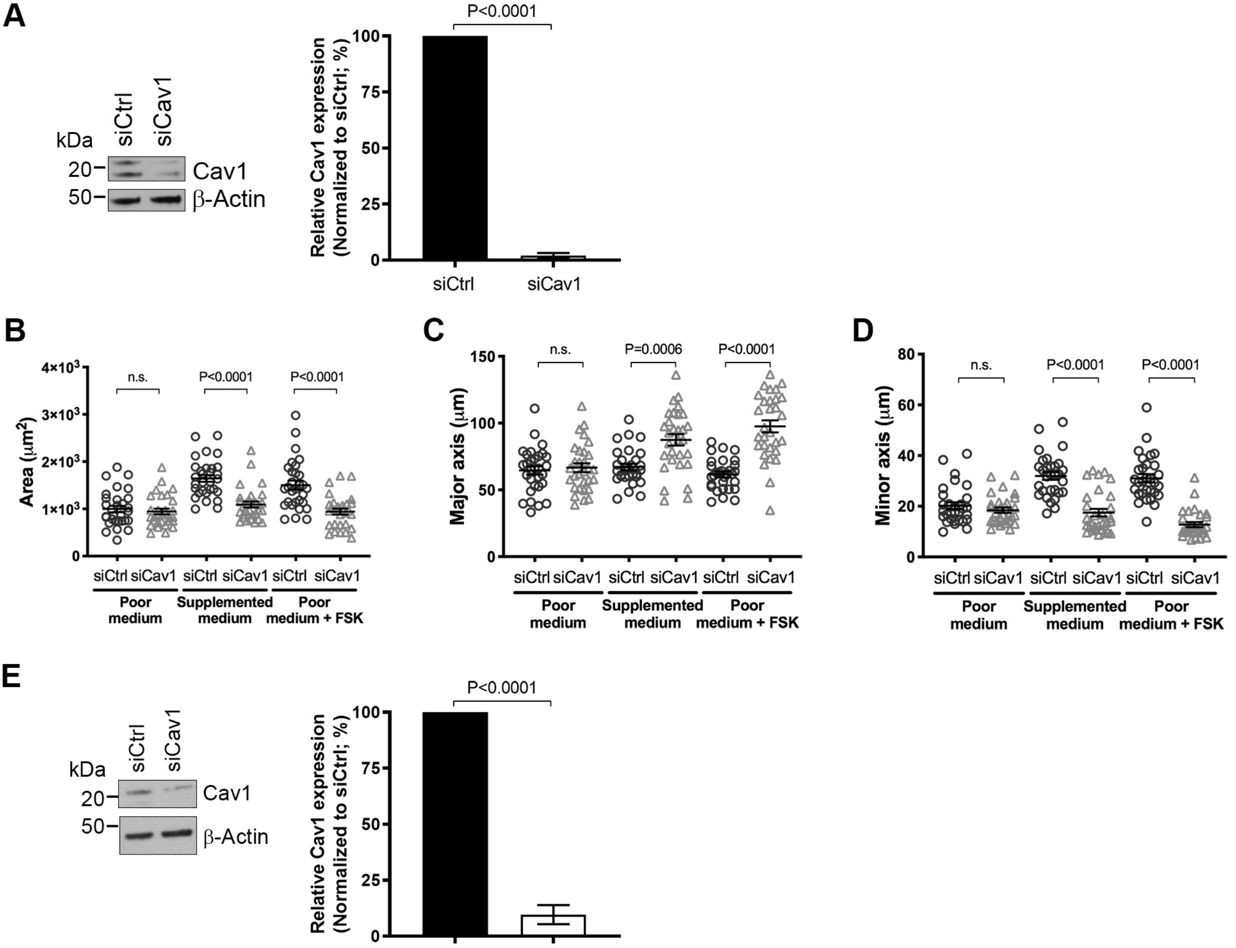

### Supplemental Figure 4

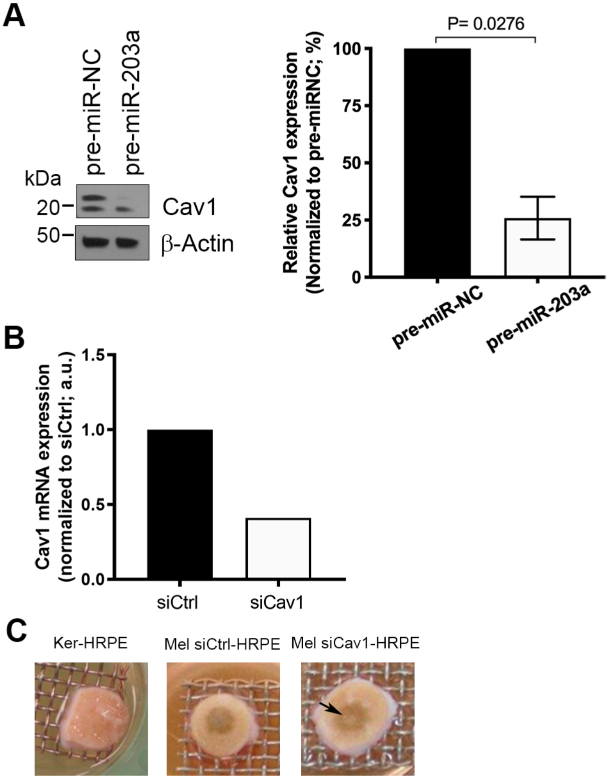
